# Can near-infrared exposure help mitigate pesticide-impaired thermoregulation in adult bees?

**DOI:** 10.64898/2026.01.07.698120

**Authors:** Tianqi Zhu, Peter Graystock, Richard J. Gill

## Abstract

For terrestrial animals, maintaining body temperature is vital for successful movement, feeding and reproduction. Exposure to environmental toxins such as pesticides, however, may place individual thermoregulation at risk. Identifying potential mitigative solutions is thus important, especially given widespread pesticide applications across landscapes are forecast to continue. In mammals, near infrared (NIR) light has been put forward as a method for countering the effects of toxin exposure, but limited evidence has been found for insects. Here, we orally expose an important insect pollinator group - adult bumblebees - to a cholinergic pesticide (a neonicotinoid) to first quantify the baseline impact on body temperature (using thermal imaging), and second to investigate how exposure to near infrared (NIR) light mediates this effect. For bumblebees kept at 18°C ambient we show, relative to control bees who maintained their body temperature at ca.30°C, that neonicotinoid exposed bees exhibited a steep (by nearly 8°C) body temperature drop over the 73hr assay showing a pattern of convergence towards ambient. We did however find some evidence that NIR can partially counter such neonicotinoid impairment. Of two different wavelengths (600nm, 850nm) and three exposure durations (5, 10 or 20 minutes per hour), we found evidence that 660 nm for 5 minutes appeared to more than halve the rate to which the neonicotinoid decreased body temperature. Yet, intriguingly, the mitigation effect of 660 nm NIR appeared to lessen as exposure time increased. We further found tentative evidence that 850 nm for 20 mins may alleviate the decline in body temperature (by nearly 40%). In conclusion, our study further highlights a mechanism by which pesticides threaten the critical functioning of non-target organisms but also provides a potential therapy worthy of further investigation.

## 1. INTRODUCTION

Pesticides have played an integral role in boosting global food production through crop protection [1], but widespread application has inadvertently placed non-target organisms at risk [2]. For the ecologically and economically important insect pollinators [3] we know that individuals can be frequently exposed to pesticides across landscapes [4–6]. Moreover, exposure to even sub-lethal levels of pesticides can have dramatic effects on individual behaviour and fitness proxies [7–13]. Yet, despite these risks to a group of animals we depend so highly on, the complexities of reaching food security targets means that increased pesticide applications are likely to continue [14]. It is, therefore, essential to improve our understanding of precisely how pesticides can impact key insect pollinator functions [15–17] if we are to develop strategies that can help mitigate acute and chronic effects in the wild [18,19].

Physiological and behavioural based thermoregulation plays a central role in determining locomotion and, in many cases, brood rearing for insect pollinators [20,21], but may be a function at risk from pesticides in the field [22,23]. For the heterothermic bumblebees, the need for individuals to warm their bodies is achieved by vibrating flight muscles to produce heat (thermogenesis) requiring significant energetic expenditure [24]. If pesticides act to interfere with thermogenesis, even if subtly, impacts on bumblebee foraging performance [25–27] and reproduction [28] are likely. For instance, we know that cholinergic pesticides (neurotoxins that target acetylcholinesterase receptors) have been reported to affect neuronal mitochondrial function and the expression of genes involved with energetic metabolism pathways [29–33]. Furthermore, such exposure has been associated with negative effects on individual thermogenesis [23] seemingly affecting the ability of individuals to thermoregulate the nest [13]. Given bees are forecast to experience greater temperature challenges under climate change and continued exposure to cholinergic pesticides (primarily through contaminated pollen and nectar [34–36], further quantification of the thermoregulatory impacts from cholinergic pesticide exposure has never been more important. It is, however, equally important to explore countermeasures.

A potential mitigative method could be the use of near-infrared radiation (NIR) therapy, exposing individuals to spectral wavelengths between 600-1000 nm. For instance, *in vitro* studies on NIR have indicated enhanced mitochondrial function and delayed cell apoptosis in isolated mammalian cells or tissues [37–39]. Moreover, *in vivo* studies on rodents indicates that NIR can reduce inflammation, accelerate wound healing, and improve outcomes of ischemic and neurological injuries [40–42]. Comparative studies on insects are limited, but an *in vivo* study on the fruit fly *Drosophila melanogaster* did show that 670 NIR exposure could increase cellular ATP levels and extended individual lifespan [43,44]. Furthermore, bumblebees exposed to a neonicotinoid pesticide show evidence of recovery in mitochondrial and motor activity for individuals also exposed to 670 nm NIR, suggested to stem from absorption of NIR by cytochrome c oxidase in mitochondrial respiration [45,46]. But whether NIR exposure to insect individuals (*in vivo*) can counter the impacts of pesticides on body temperature and thermoregulation has, to our knowledge, not been investigated.

Here, we look to accurately quantify how exposure of individual bumblebees (*Bombus terrestris audax*) to a neonicotinoid pesticide (imidacloprid) impacts the ability of individual workers to maintain body temperature. We then test whether NIR exposure can help mitigate any impact (Fig. 1). We investigated this by measuring thorax temperature through thermal imaging after exposing imidacloprid-treated bumblebee adult workers to 660 or 850 nm wavelengths each for three different rates of exposure (5, 10 and 20 mins per hour). We tested the hypothesis that NIR therapy would allow pesticide exposed bumblebees to partially or fully restore their thorax temperature.

**Figure 1.**
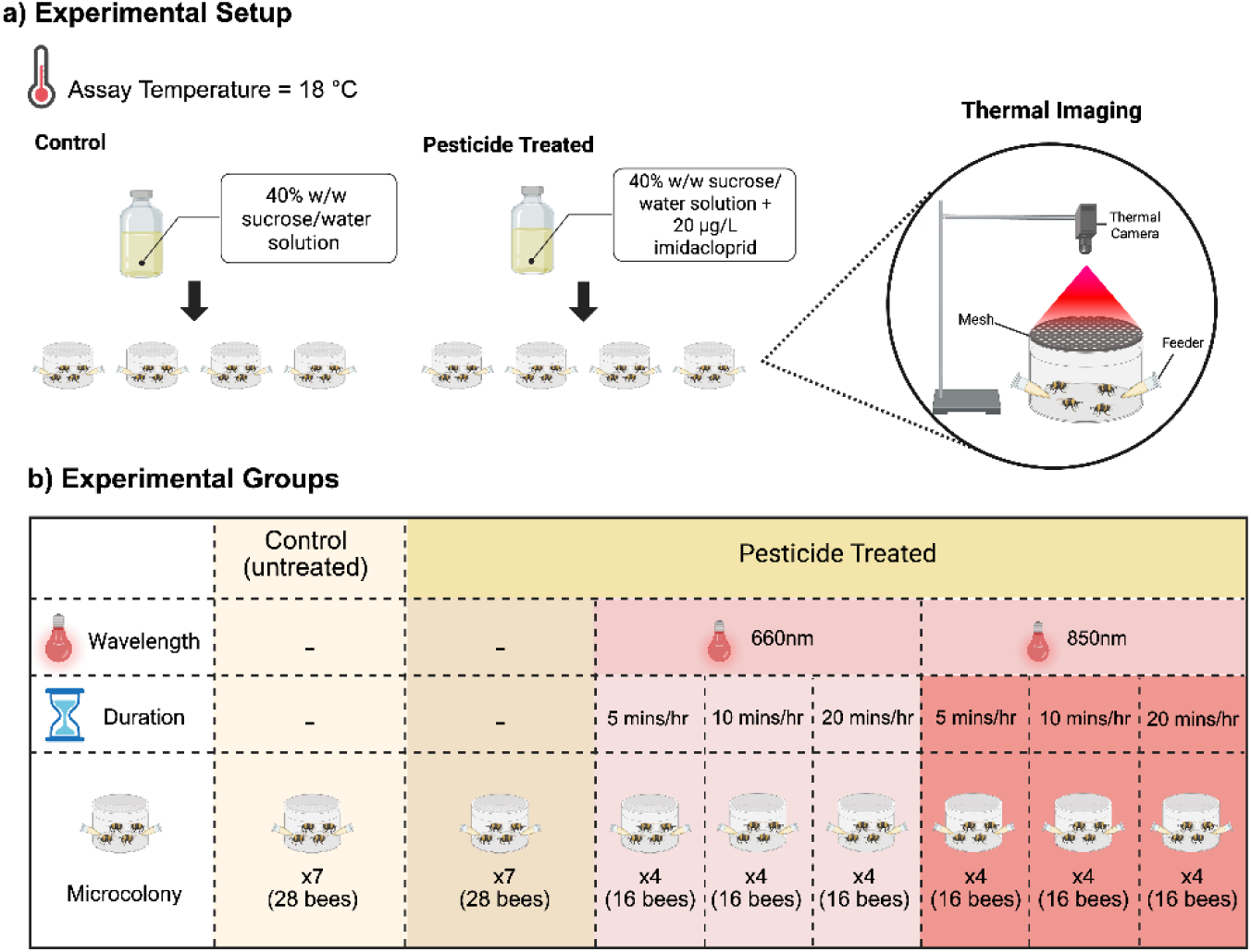
Experimental design testing the effect of NIR on mitigating the impact of pesticide (imidacloprid) exposed bees. **a)** Illustration of the experimental setup; **b)** The factorial design of our treatments with sample sizes shown. Bumblebee microcolonies (four individual bees per microcolony) were assigned either to a control treatment (no pesticide, no NIR), pesticide only (with pesticide, no NIR), and NIR-treated group with pesticide at 660nm or 850 nm for 0, 5, 10, 20 min/h. Bees were kept in microcolonies covered with a metal mesh and fed via Eppendorf tube feeders. Thorax temperature was monitored using a thermal imaging camera, which recorded the temperature at the same position above each microcolony. The experiment was conducted at a standardised ambient temperature of 18 °C.

## 2. Methods

### 2.1 Bee setups and route of pesticide exposure

Two bumblebee (*B. terrestris audax*) ‘parental colonies’ were obtained from the commercial supplier Biobest (distributor: Agralan Ltd.). Upon delivery, they were housed in a dark controlled environment (CE) room, maintained at a constant 18°C and 60% relative humidity (RH). Any sugar reservoirs that came with the colonies were removed and replaced with gravity feeders providing a 40% w/w sucrose/water solution *ad libitum*, with pollen (four grams) provided every three days inside a 45 mm diameter petri dish. After two weeks, 76 worker bees per parental colony (total = 152) were randomly removed and placed in cohorts of four sisters per transparent plastic container (upper and lower diameter = 11.5 and 9.0 cm; height = 5.5 cm) to produce a set of 38 microcolonies. The roof of each microcolony container was constructed from a plastic 3D printed ring with aluminium wire mesh (hole size = 4 mm), which allowed airflow and thermal imaging of individuals whilst containing bees throughout the assay. Microcolonies were fed *ad libitum* with untreated 40% sucrose solution and left for 24 hours before the 4-day assay began. The assay was run under continuous 18°C at 60% RH, with two independent but standardised replications of this experiment (using different worker individuals) conducted consecutively. The room remained under darkness when the assay started to ensure NIR was the only light exposure.

Two holes were made in the side walls of the containers in which 1.5mL Eppendorf tubes could be inserted, each with a small hole (2-3mm in diameter) at the bottom for the bee to access using their tongue. For pesticide treatment microcolonies, feeders contained 40% sucrose solutions spiked with 20 µg/L imidacloprid, whereas for control microcolonies, feeders were filled with untreated sucrose solution. At the start of the assay, we provisioned a standardised 1.5 ml of sucrose solution per feeder (total = 3mL) to each microcolony and replaced it every day (before the first thorax temperature measurement; see below), and daily food consumption was recorded by measuring any remaining sucrose solution on replacement.

### 2.2 Pesticide and NIR exposure treatments

Each of the 38 microcolonies was first assigned to one of three treatment categories: control (untreated sucrose solution; n = 7 microcolonies, 28 workers), pesticide-only (sucrose solution with imidacloprid; n = 7 microcolonies, 28 workers), and pesticide + NIR (n = 24 microcolonies, 96 workers). Then, for the pesticide + NIR treatment group, we further divided microcolonies into receiving different wavelengths using either a 660 nm LED bulb (7W) or 850 nm LED bulb (12W) each with 60° lenses. Thes two wavelength treatments were combined with different exposure durations (5, 10, or 20 min/hr), with four microcolonies (n = 16 workers) per combinatorial treatment (Fig. 1). The two different wavelengths were chosen because they have been commonly used in other medical and biological applications [44,45,47,48]. Bulbs were attached to multiple independent desk lamps, with each placed directly above a microcolony and enclosed in a fashioned box when exposed to prevent cross-exposure between NIR treatments. NIR exposure was given as periodic doses per hour; for example, for the ‘*pesticide +5*’ treatment group, individuals would be exposed to NIR for 5 minutes followed by 55 minutes without exposure, which was repeated hourly. This was conducted between the hours of 12:00 and 18:00. NIR treatment groups had their exposure initiated in a staggered manner, allowing us to expose all colonies in a standardised way with sufficient time to capture replicate thermal images throughout the day.

### 2.3 Thermal imaging

Repeat thermal imaging of the bees was taken every hour between 12:00 to 17:00 to give six measures per worker per day for assay days 1-3 (total = 18 measures), and one last measure taken at ∼12:00 on day four. We used a FLIR Pro camera (emissivity set at 0.95) with thermal images analysed using the smart phone application FLIR Tools (Version 2.11.1, Atlas Mobile SDK 2.9.1, © 2015, FLIR Systems, Inc.) and the FLIR Thermal Studio software (Version 2.0.45.0, Atlas SDK 7.5.1.1955, released on June 2, 2025), which transfers thermal pixels into digital temperature data. Thorax temperature was determined by selecting the pixel with the highest temperature within the thoracic region of each bee from the thermal images. Calibration of the thermal camera was performed using an ‘artificial bee’ which was a polystyrene cube (dimensions = 30 x 30 × 10 mm) covered entirely with black electrical tape with the same emissivity as our studied bumblebees (ε = 0.95 ± 0.05). The surface temperature of the black cube would reflect that of the environmental temperature around the microcolony, allowing us to calculate and distinguish the effects of individual thermogenesis from bulb heat on thorax temperature across treatments (Fig. S1).

### 2.4 Statistical analysis

Statistical analyses were conducted using R software (version 4.4.2, R Core Team 2024) with graphs made using ggplot2 [49]. Two linear mixed effects models (LMMs) were fitted to the data. Model 1 looked to predict thorax temperature as a function of *pesticide* exposure and its interaction with *time*, with control as the intercept for comparison (Control vs. Imidacloprid). Model 2 looked to predict thorax temperature as a function of NIR treatments with different wavelengths (660nm and 850nm) and exposure duration (5, 10, 20 min), resulting in seven levels as fixed factors and their interaction with *time.* For model 2, we assigned the ‘Imidacloprid without NIR’ as the intercept for comparison. For both models, we accounted for non-independence between workers from the same microcolony by considering *container* as a random effect, and incorporated food *consumption* per bee as a covariate, given that we know metabolic heat production is dependent on carbohydrate intake. To estimate food consumption per bee, we divided the daily food intake (mg) of each microcolony by the number of workers within that microcolony (i.e. four bees).

With the assay being run over four days, our temporal thorax temperature data inevitably had temporal data gaps (19hrs between days) which led to assumption violations when fitting the linear models. Thus, for the purpose of broadly testing chronic effects and simplifying our statistical analysis, we modelled each repeated thorax temperature as chronological order (integer) of measurements (*timeseq* = 1-19; but for models fitted using raw time please see Tables S1-S2). The assumptions of each model were then checked again using the performance package [50], and the residuals of both models were approximately normally distributed. Our data does exhibit mild heteroscedasticity, but some of the uneven variance was absorbed by the random-effect structure. We further mean-centered the time variable due to multicollinearity between NIR treatment and its interaction with time (VIF > 30-100), which dropped the VIF values to <5 and significantly improved model stability. Meanwhile, the estimated slope remained unchanged, indicating that mean-centring did not affect the ecological interpretation of the model.

## 3. RESULTS

### 3.1 Effect of pesticide exposure on thorax temperature

Control bees maintained a stable thorax surface temperature throughout the assay, with a mean model estimate of 29.7°C (95% CI: 28.8–30.6°C; Fig. 2). In contrast, pesticide-treated bees were unable to maintain their thorax temperature, exhibiting a consistent decline with time (LMM: *pesticide x time*: t = −8.21, p < 0.001; Table S3). For pesticide-treated bees, the estimated rate of decrease per sequential thorax measure was −0.417°C (95% CI: –0.488 to –0.345°C), which equated to an estimated fall of mean thorax temperature in pesticide-treated bees to 23.8°C (95% CI: 22.6–25.1°C) by the end of the 73hr assay. Importantly, food consumption also independently mediated bumblebee thorax temperature, with a decrease of 0.0095°C for each 1 mg less of sucrose consumed (t = 7.14, p < 0.001).

**Figure 2.**
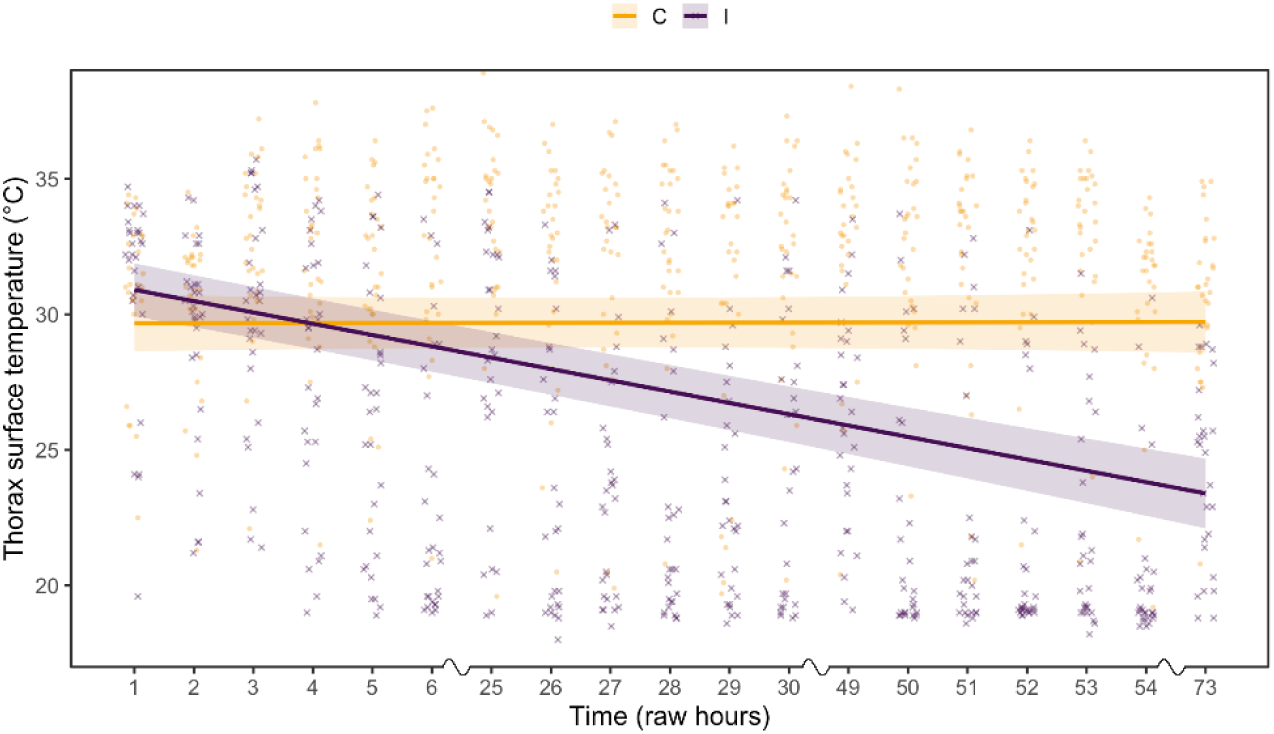
Repeat measures of bumblebee thorax temperature over time for control bees (provisioned untreated nectar substitute throughout the assay; n=28 individuals; yellow) and pesticide-exposed bees (provisioned imidacloprid-spiked nectar substitute; n=28 individuals; purple). Background data points represent raw measurements per individual, with fitted lines and shaded ribbons showing the LLM estimated marginal means and 95% confidence intervals, respectively. *Note:* the LLM estimates are adjusted to account for the fixed effect of consumption and random effect of microcolony leading to estimates looking slightly lower or higher relative to the raw data cloud. The LLM model also considered time as a chronological order of 19 repeat measurements, but for visual purposes the hours since the assay started are labelled along the x-axis.

### 3.2 Does NIR counter pesticide impacts on thorax temperature

At the beginning of the assay, the thorax temperatures of the bumblebees showed no significant difference between the pesticide-treated-only and NIR + pesticide-treated groups (LMM: all p>0.4). The estimated delineation rate of temperature per sequential thorax measure for the pesticide-treated-only bees was −0.427 °C (95% CI: –0.504 to –0.350) which was close to the estimate from model 1. We did, however, find subtle evidence that NIR can slow the rate of decline in thoracic surface temperature in pesticide-exposed bumblebees (Fig. 3). Among all the NIR *wavelength* and *exposure duration* combinations, the most effective treatment was 5-minute exposure at 660nm (t = 5.05, p < 0.001; Table S4), with a shallower delineation rate per sequential thorax measure of −0.107 °C (95% CI: −0.208 to −0.006). Additionally, 10-minute exposure at 660nm and 20-minute 660nm treatment group also showed significant alleviation of the drop in the thorax temperature of bees (t=4.12, p<0.001; t=2.02, p=0.043), with an estimated delineation rate per sequential thorax measure of −0.163 °C (95% CI: –0.208 to –0.006) and - 0.299 °C (95% CI: –0.405 to –0.192). The alleviation effect of the treatment groups using 660nm NIR decreased as the NIR exposure time increased.

**Figure 3.**
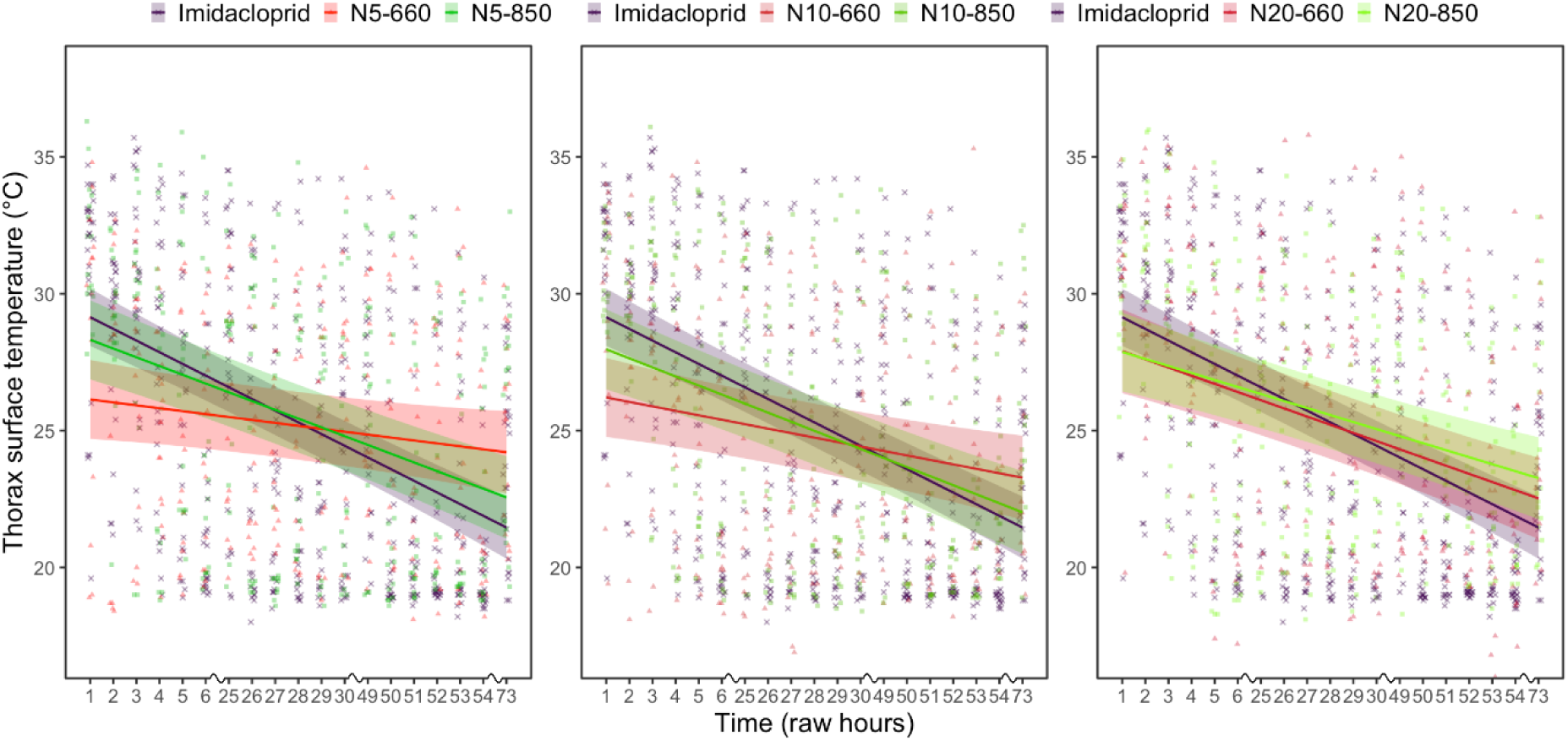
Repeat measures of bumblebee thorax temperature over time for the pesticide-exposed bees that received NIR (red = 660 nm; green = 850nm; total = 124 individuals) relative to the pesticide-exposed bees that received no NIR (purple). The left panel shows a comparison of bees exposed for 5 minutes per hour for each wavelength, the middle panel for 10 minutes per hour and the right panel for 20 minutes per hour. Background data points represent raw measurements per individual, with the fitted lines and shaded ribbons showing the LLM estimated marginal means and 95% confidence intervals, respectively. *Note:* the LLM model accounted for consumption and considered time as a chronological order of 19 repeat measurements, but for visual purposes the hours since the assay started are labelled along the x-axis.

In contrast, bees with shorter exposure duration (5 and 10 minutes) to 850nm NIR showed no difference compared with pesticide-only treated bees (t=1.70, p=0.089; t=1.52, p=0.128). However, bees treated with 20-minute 850nm NIR exposure showed a shallower decline in thorax temperature (t=2.72, p=0.006), with an estimated delineation rate per sequential thorax measure of −0.256°C (95% CI: −0.357 to −0.155). Again, we found that higher food consumption significantly increased thorax temperature (t = 6.14, p < 0.05).

## 4. DISCUSSION

Here, we quantify the rate to which chronic exposure to a cholinergic pesticide (the neonicotinoid, imidacloprid) in adult bumblebee workers significantly reduces thorax temperature. We show that after 73hrs the body temperature of pesticide exposed bumblebee workers was converging towards the ambient temperature, being around 20% lower than control bees which maintained their body temperature >10°C above the ambient. We then provide evidence that simultaneous exposure to NIR may be able to partially counter some of this impact, although thorax temperature by the end of the assay was still below that of control bees. We further reiterate the importance of diet, in which we quantify how consumption rate of the nectar substitute (sucrose solution) contributes to raising body temperature.

### i) Neonicotinoid exposure impairs the regulation of bumblebee body temperature

Our findings align with others showing that neonicotinoid pesticide exposure can negatively affect the thermoregulation abilities of bees [22]. Determining the physiological mechanisms being impacted requires further investigation, but three non-mutually exclusive explanations come to mind. 1) Pesticide residues within the body are directly interfering with metabolic pathways, rendering bee’s physically incapable of raising their body temperature. Thorax temperature is an essential indicator of metabolic function [51,52], and neonicotinoids have been shown to interfere with oxidative phosphorylation (the formation of ATP) and respiration [53] and thus likely impairing the role of flight muscles in thermogenesis. Additionally, studies have linked neonicotinoid exposure to reduced locomotion in bumblebees [25,54], which, in turn, could lead to a negative feedback loop of drops in body temperature. 2) Acute pesticide exposure causes sensory impairment and reduced foraging motivation [55,56], stemming from secondary consequences of neurotoxic disruption and thus altered activity patterns. Thus, even if bees have the physical potential to raise their body temperature, perhaps they are not motivated to do so. 3) Thermogenesis in bees is a highly energy-demanding activity that requires rapid carbohydrate intake [57]. Yet our findings, like others, show reduced food consumption in bumblebees after neonicotinoid exposure suggesting a reduced appetite [16,58]. Without a suitable carbohydrate intake, bees may be unable to raise their temperature due to a depauperate energy source. Indeed, we found a 0.095°C decrease in thorax temperature for each 10 mg less of 40% sucrose solution consumed, and a previous study showed that for each 1 mol/L sucrose concentration increment provisioned to bumblebees (*Bombus wilmattae*) increase thorax temperature by 4.2°C [59]. Furthermore, with other studies indicating that nutritional stress can exacerbate pesticide impacts to bees [60], the decrease in thorax temperature likely results from a combination of neurotoxic damage and nutritional shortage. Of course, the three explanations above are non-mutually exclusive, and further investigation is needed to identify and distinguish the molecular, physiological, and neurological mechanisms being affected.

### ii) Tentative evidence that NIR exposure to adult bees can partially counter pesticide impacts

Previous findings suggest that NIR can positively affect both vertebrates and invertebrates by enhancing mitochondrial function and increasing ATP production, thereby improving physiological performance [37–39,43,44]. Our findings suggest that NIR, whilst not fully mitigating the impacts of the pesticide, did show evidence of a subtle counter-effect. As our results show, the rate of decline in bumblebee body temperature was, to some extent, alleviated in most NIR treatment groups. Powner *et al.* (2021) reported that just 1min of NIR treatment can begin to correct respiratory dysfunction in bumblebees caused by imidacloprid exposure, and that extending the exposure time does not further increase the benefits. In our study, bumblebees were treated with NIR for at least 5 minutes per hour, and similar mitigation effects were observed. Research also suggests that intensified and repeated NIR exposure might induce oxidative stress and other cellular responses that counteract some of the effects of NIR [45,46]. In other words, our treatments may have exceeded the optimal NIR level, leading to saturation of photobiological stimulation. Indeed the 5-minute exposure compared with the 10- and 20-minute treatments appeared to provide the greatest beneficial effect.

Whilst we provide some indicative findings, more work is needed to elucidate the underlying mechanisms of NIR therapy in bees. For example, measures of a pesticide and NIR interactive effect on ATP levels, metabolic and neuronal activity could be looked at and potentially over longer observation period to determine an acute versus chronic therapy. It would also be worth determining the degree to which NIR is needed to penetrate internal tissues (such as flight muscles) of individual bees. Adult bumblebee workers possess a hard and thick exoskeleton, which might limit the NIR penetration to the cellular-level processes where the functional responses occur. Most studies to date have examined direct exposure of NIR to cells *in vitro* without little to no barrier to radiation [38,42]. With bee larva lacking a hard cuticle, an interesting line of investigation may be to test the effects of deeper NIR penetration in larva that are being reared in a pesticide exposure environment (i.e. simulating residues within the colony).

### iii) Conclusion

The impacts that the tested cholinergic pesticide can have on worker body temperature is likely to have serious implications for individual task performance and colony success in the field [27]. For bumblebees, flight is energetically expensive because individuals must warm their flight muscles to >30 °C for activation [59]. The rapid decline in thorax temperature shown in this study helps explain the growing body of evidence linking pesticides to reduced foraging range [25] and diminished pollen collection [7,26]. Moreover, a recent study showed that pesticide induced impairment to individual bumblebee thermoregulation can translate to impacts on pupal development [28]. Overall, our findings provide further evidence that impairment of the key function, body thermoregulation, may have contributed to why bees have been shown to perform poorly in pesticide exposure landscapes [61–65]. Whilst further investigation is required, the application of NIR in commercial settings, such as adapting managed hives with fitted NIR LEDs, could represent a potential mitigative method for helping bees to buffer the effects of pesticide exposure. Its use would seem especially important during any later-season cold snaps that are becoming more frequent under climate change.

## Contributions

RJG and PG conceived the project; TZ and RJG designed and set up the experiment; TZ collected and analysed the data; TZ and RJG wrote the paper with input from PG.

## Acknowledgements

We are grateful to Owen Stewart for providing feedback on the manuscript. Thanks also go to Paul Beasley, Morgan Berrell, and Annabel West for technical support, and to Kieran Storer and Yao Yao for piloting the experimental setups.

## Supplementary Tables

**Table S1.** Linear mixed-effects model using raw experimental time. Model: Temperature ∼ Group * time + Consumptionperbee + (1|Container)

| TERM | ESTIMATE |  | STD. ERROR | DF | T VALUE | P-VALUE |
| --- | --- | --- | --- | --- | --- | --- |
| (INTERCEPT) | 25.1239 |  | 0.9575 | 29.776 | 26.240 | < 2e-16 *** |
| GROUPI | 0.5287 |  | 0.7501 | 26.066 | 0.705 | 0.487 |
| TIME | 0.0026 |  | 0.0094 | 993.418 | 0.272 | 0.785 |
| CONSUMPTIONPERBEE | 0.0099 |  | 0.0013 | 26.398 | 7.439 | 6.11e-08 *** |
| GROUPI:TIME | -0.0996 |  | 0.0137 | 855.628 | -6.993 | 5.43e-12 *** |
| RANDOM EFFECT | Variance | Std. Dev. | Groups |  |  |  |
| CONTAINER | 0.1688 | 0.4109 | 8 levels |  |  |  |
| RESIDUAL | 16.6424 | 4.0795 | — |  |  |  |

**Table S2.**
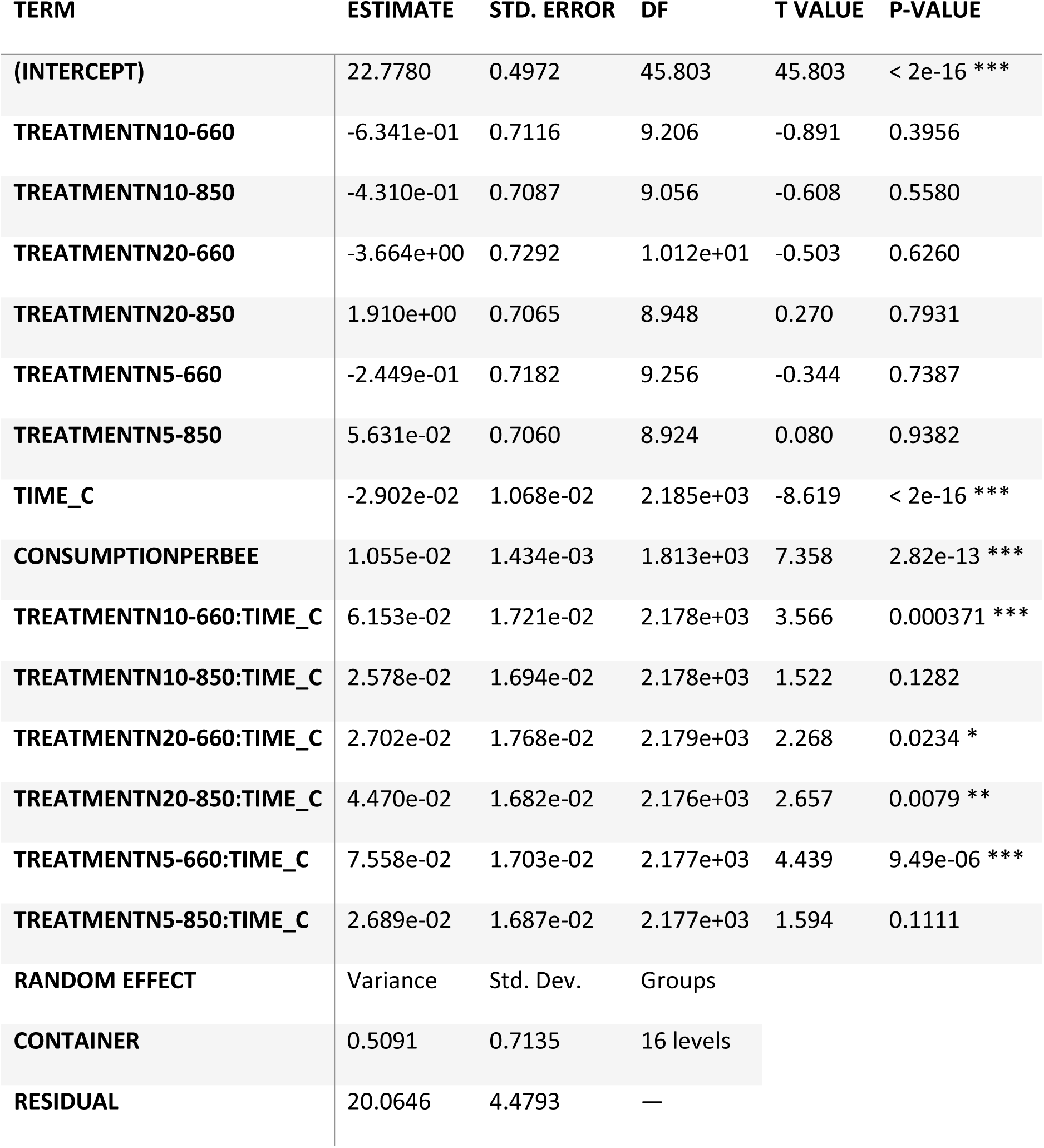
Linear mixed-effects model using raw experimental time. Model: Temperature ∼ Treatment * time_c + Consumptionperbee + (1 | Container)

| TERM | ESTIMATE | STD. ERROR | DF | T VALUE | P-VALUE |
| --- | --- | --- | --- | --- | --- |
| (INTERCEPT) | 22.7780 | 0.4972 | 45.803 | 45.803 | < 2e-16 *** |
| TREATMENTN10-660 | -6.341e-01 | 0.7116 | 9.206 | -0.891 | 0.3956 |
| TREATMENTN10-850 | -4.310e-01 | 0.7087 | 9.056 | -0.608 | 0.5580 |
| TREATMENTN20-660 | -3.664e+00 | 0.7292 | 1.012e+01 | -0.503 | 0.6260 |
| TREATMENTN20-850 | 1.910e+00 | 0.7065 | 8.948 | 0.270 | 0.7931 |
| TREATMENTN5-660 | -2.449e-01 | 0.7182 | 9.256 | -0.344 | 0.7387 |
| TREATMENTN5-850 | 5.631e-02 | 0.7060 | 8.924 | 0.080 | 0.9382 |
| TIME_C | -2.902e-02 | 1.068e-02 | 2.185e+03 | -8.619 | < 2e-16 *** |
| CONSUMPTIONPERBEE | 1.055e-02 | 1.434e-03 | 1.813e+03 | 7.358 | 2.82e-13 *** |
| TREATMENTN10-660:TIME_C | 6.153e-02 | 1.721e-02 | 2.178e+03 | 3.566 | 0.000371 *** |
| TREATMENTN10-850:TIME_C | 2.578e-02 | 1.694e-02 | 2.178e+03 | 1.522 | 0.1282 |
| TREATMENTN20-660:TIME_C | 2.702e-02 | 1.768e-02 | 2.179e+03 | 2.268 | 0.0234 * |
| TREATMENTN20-850:TIME_C | 4.470e-02 | 1.682e-02 | 2.176e+03 | 2.657 | 0.0079 ** |
| TREATMENTN5-660:TIME_C | 7.558e-02 | 1.703e-02 | 2.177e+03 | 4.439 | 9.49e-06 *** |
| TREATMENTN5-850:TIME_C | 2.689e-02 | 1.687e-02 | 2.177e+03 | 1.594 | 0.1111 |
| RANDOM EFFECT | Variance | Std. Dev. | Groups |  |  |
| CONTAINER | 0.5091 | 0.7135 | 16 levels |  |  |
| RESIDUAL | 20.0646 | 4.4793 | — |  |  |

**Table S3.**
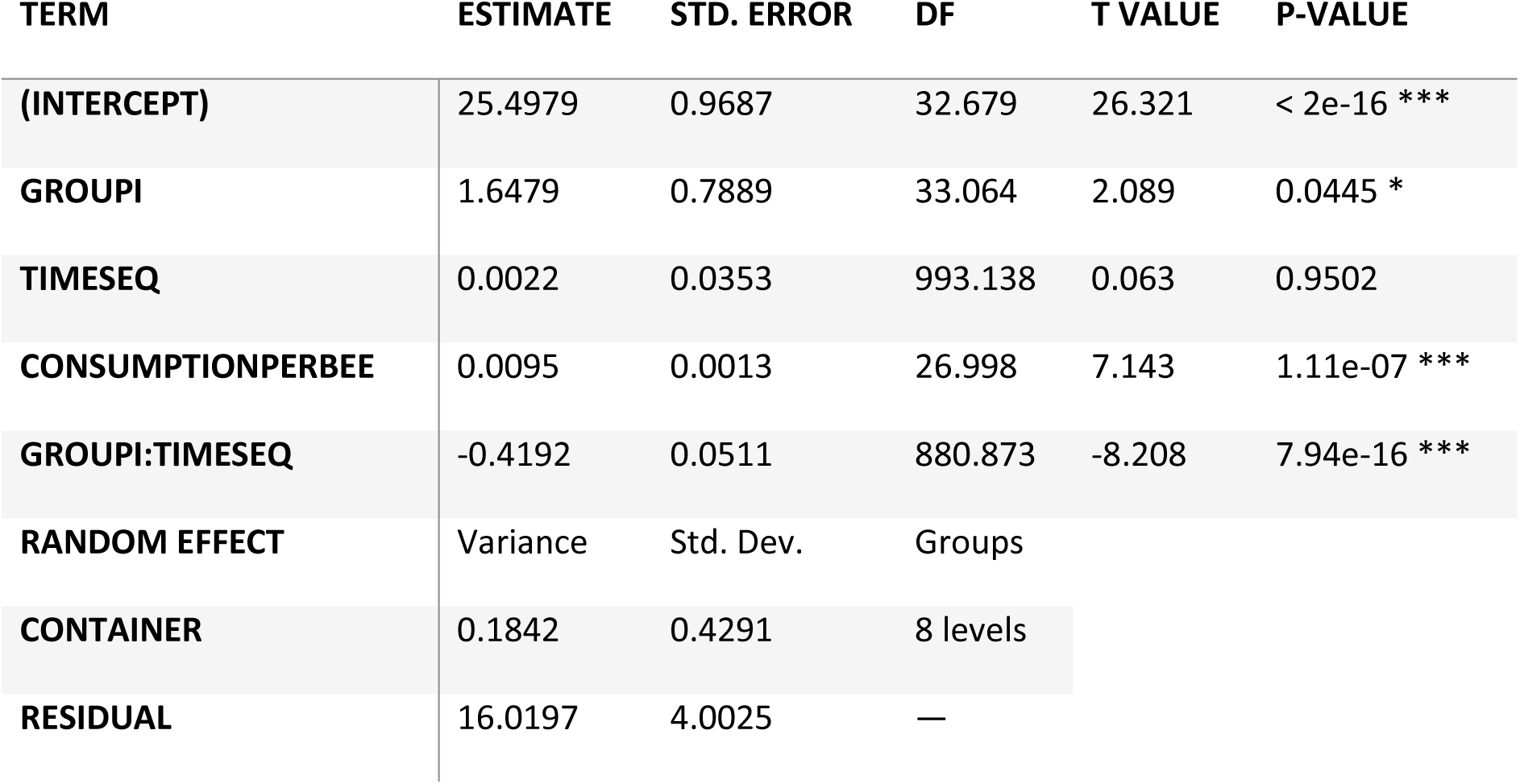
Linear mixed-effects model using integer time index (Main model 1) Model: Temperature ∼ Group * timeseq + Consumptionperbee + (1|Container)

**Table S4.** Linear mixed-effects model using centered integer time index (Main model 2. ) Model: Temperature ∼ Treatment * timeseq_c + Consumptionperbee + (1 | Container)

| TERM |  | ESTIMATE | STD. ERROR | DF | T VALUE | P-VALUE |
| --- | --- | --- | --- | --- | --- | --- |
| (INTERCEPT) |  | 23.22 | 0.4737 | 49.027 | 49.027 | < 2e-16 *** |
| TREATMENTN10-660 |  | -0.5554 | 0.6778 | 9.156 | -0.819 | 0.4330 |
| TREATMENTN10-850 |  | -0.3077 | 0.6750 | 9.006 | -0.456 | 0.6593 |
| TREATMENTN20-660 |  | -0.8730 | 0.6945 | 10.007 | -0.125 | 0.9027 |
| TREATMENTN20-850 |  | 0.2669 | 0.6728 | 8.894 | 0.397 | 0.7009 |
| TREATMENTN5-660 |  | -1.2626 | 0.6786 | 9.197 | -0.186 | 0.8564 |
| TREATMENTN5-850 |  | 0.1376 | 0.6724 | 8.872 | 0.205 | 0.8424 |
| TIMESEQ_C |  | -0.4270 | 0.0395 | 2184.0 | -10.822 | < 2e-16 *** |
| CONSUMPTIONPERBEE |  | 0.0223 | 0.0032 | 1822.0 | 6.143 | 9.91e-10 *** |
| TREATMENTN10-660:TIMESEQ_C |  | 0.2640 | 0.0641 | 2178.0 | 4.119 | 3.94e-05 *** |
| TREATMENTN10-850:TIMESEQ_C |  | 0.1594 | 0.0659 | 2177.0 | 1.523 | 0.1279 |
| TREATMENTN20-660:TIMESEQ_C |  | 0.1281 | 0.0634 | 2179.0 | 2.021 | 0.0434 * |
| TREATMENTN20-850:TIMESEQ_C |  | 0.1707 | 0.0626 | 2176.0 | 2.724 | 0.0064 ** |
| TREATMENTN5-660:TIMESEQ_C |  | 0.5138 | 0.0640 | 2177.0 | 5.045 | 4.91e-07 *** |
| TREATMENTN5-850:TIMESEQ_C |  | 0.1069 | 0.0628 | 2177.0 | 1.703 | 0.0887 |
| RANDOM EFFECT |  | Variance | Std. Dev. | Groups |  |  |
| CONTAINER |  | 0.4538 | 0.6737 | 16 levels |  |  |
| RESIDUAL |  | 19.3527 | 4.3992 | — |  |  |

## Supplementary Figures

**Figure S1.**
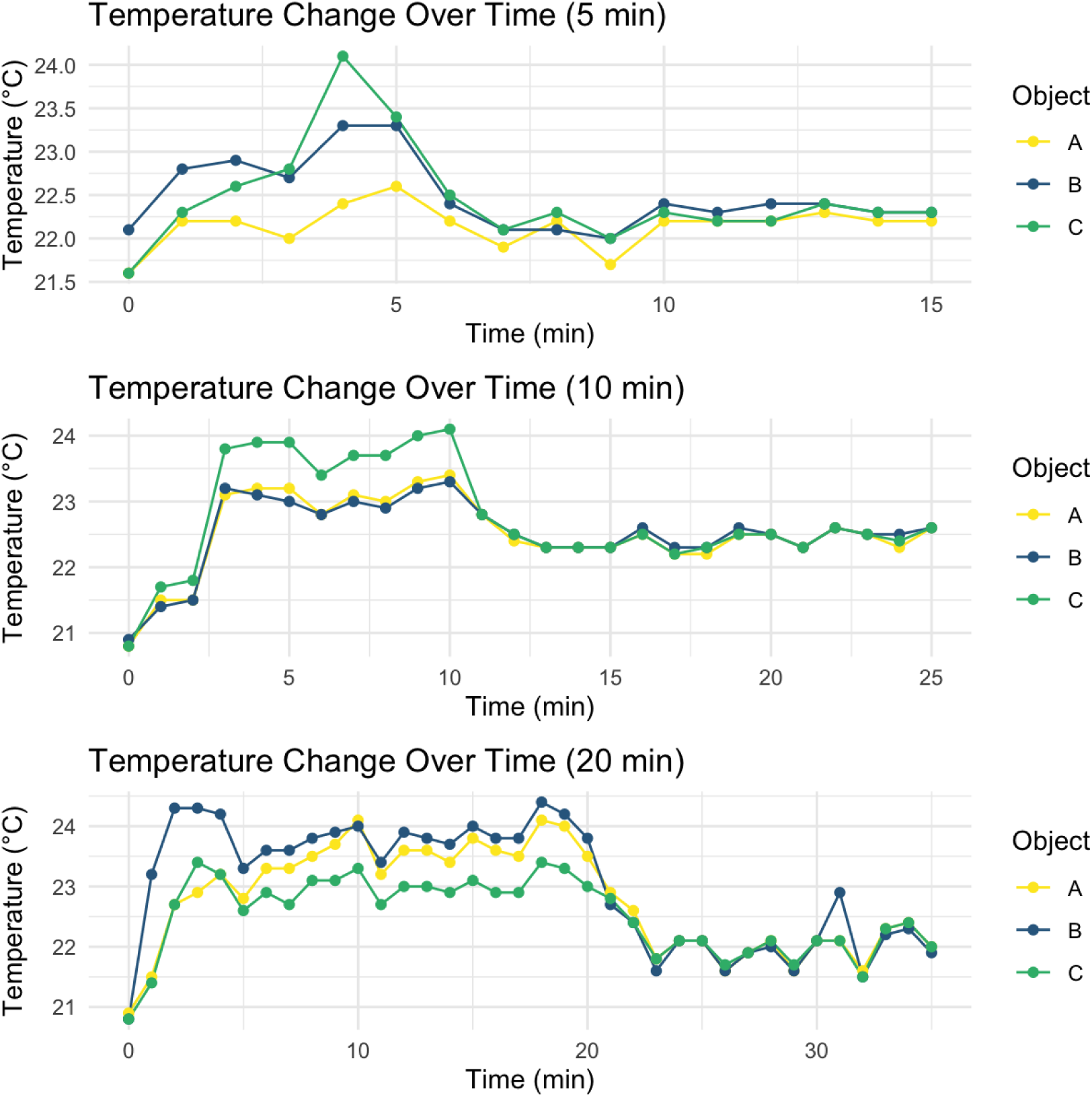
Temperature over time for the black cube references used for camera calibration of thermal emissivity. Three cubes (A, B and C) were placed in a 21°C room exposed to NIR for 5 minutes (top) then separately for 10 minutes (middle) and again separately for 20 minutes (bottom) to determine if any heat-up effect from the NIR bulb irradiation occurs. The result shows that NIR bulbs can potentially heat bees to around 2°C (Figure S2). Thus, to account for the heat-up effect (i.e. calibrate), a black cube was placed next to each microcolony when being exposed to NIR with the temperature difference (based off the thermal image). Taking the temperature difference between the black cube and the 18°C ambient, we could subtract this from the measured bee temperature.

## REFERENCES

1. Popp J, Pető K, Nagy J. 2012 Pesticide productivity and food security. A review. Agronomy for Sustainable Development 33:1 33, 243–255. (doi:10.1007/S13593-012-0105-X)

2. Wan NF et al. 2025 Pesticides have negative effects on non-target organisms. Nature Communications 16:1 16, 1360-. (doi:10.1038/s41467-025-56732-x)

3. Potts SG et al. 2016 Safeguarding pollinators and their values to human well-being. Nature 540, 220–229. (doi:10.1038/nature20588)

4. Raine NE, Rundlöf M. 2024 Pesticide Exposure and Effects on Non-Apis Bees. Annu Rev Entomol 69, 551–576. (doi:10.1146/ANNUREV-ENTO-040323-020625/CITE/REFWORKS)

5. Gradish AE et al. 2019 Comparison of Pesticide Exposure in Honey Bees (Hymenoptera: Apidae) and Bumble Bees (Hymenoptera: Apidae): Implications for Risk Assessments. Environ Entomol 48, 12–21. (doi:10.1093/EE/NVY168)

6. Main AR, Hladik ML, Webb EB, Goyne KW, Mengel D. 2020 Beyond neonicotinoids – Wild pollinators are exposed to a range of pesticides while foraging in agroecosystems. Science of The Total Environment 742, 140436. (doi:10.1016/J.SCITOTENV.2020.140436)

7. Gill RJ, Ramos-Rodriguez O, Raine NE. 2012 Combined pesticide exposure severely affects individual- and colony-level traits in bees. Nature 491, 105–8. (doi:10.1038/nature11585)

8. Whitehorn PR, O’Connor S, Wackers FL, Goulson D. 2012 Neonicotinoid pesticide reduces bumble bee colony growth and queen production. Science 336, 351–352. (doi:10.1126/science.1215025)

9. Bryden J, Gill RJ, Mitton RAA, Raine NE, Jansen VAA. 2013 Chronic sublethal stress causes bee colony failure. Ecol Lett 16, 1463–1469. (doi:10.1111/ele.12188)

10. Siviter H, Brown MJF, Leadbeater E. 2018 Sulfoxaflor exposure reduces bumblebee reproductive success. Nature 561, 109–112. (doi:10.1038/s41586-018-0430-6)

11. Siviter H, Richman SK, Muth F. 2021 Field-realistic neonicotinoid exposure has sub-lethal effects on non-Apis bees: A meta-analysis. Ecol Lett 24, 2586–2597. (doi:10.1111/ELE.13873)

12. Chole H, De Guinea M, Woodard SH, Bloch G. 2022 Field-realistic concentrations of a neonicotinoid insecticide influence socially regulated brood development in a bumblebee. Proceedings of the Royal Society B: Biological Sciences 289. (doi:10.1098/RSPB.2022.0253/104042)

13. Crall JD et al. 2018 Neonicotinoid exposure disrupts bumblebee nest behavior, social networks, and thermoregulation. Science 362, 683–686. (doi:10.1126/science.aat1598)

14. Shattuck A, Werner M, Mempel F, Dunivin Z, Galt R. 2023 Global pesticide use and trade database (GloPUT): New estimates show pesticide use trends in low-income countries substantially underestimated. Global Environmental Change 81. (doi:10.1016/j.gloenvcha.2023.102693)

15. Stanley DA, Raine NE. 2016 Chronic exposure to a neonicotinoid pesticide alters the interactions between bumblebees and wild plants. Funct Ecol 30, 1132–1139. (doi:10.1111/1365-2435.12644)

16. Kenna D, Graystock P, Gill RJ. 2023 Toxic temperatures: bee behaviours exhibit divergent pesticide toxicity relationships with warming. Global Change Biology, 29(11), 2981–2998.

17. Chagnon M et al. 2014 Risks of large-scale use of systemic insecticides to ecosystem functioning and services. Environmental Science and Pollution Research 22:1 22, 119–134. (doi:10.1007/S11356-014-3277-X)

18. Rothman JA, Leger L, Graystock P, Russell K, McFrederick QS. 2019 The bumble bee microbiome increases survival of bees exposed to selenate toxicity. Environ Microbiol (doi:10.1111/1462-2920.14641)

19. Zhu Y et al. 2021 Estimation of Apple Flowering Frost Loss for Fruit Yield Based on Gridded Meteorological and Remote Sensing Data in Luochuan, Shaanxi Province, China. Remote Sensing, 13, 1630 13, 1630. (doi:10.3390/RS13091630)

20. Tosi S, Démares FJ, Nicolson SW, Medrzycki P, Pirk CWW, Human H. 2016 Effects of a neonicotinoid pesticide on thermoregulation of African honey bees (Apis mellifera scutellata). J Insect Physiol (doi:10.1016/j.jinsphys.2016.08.010)

21. Potts R, Clarke RM, Oldfield SE, Wood LK, Hempel de Ibarra N, Cresswell JE. 2018 The effect of dietary neonicotinoid pesticides on non-flight thermogenesis in worker bumble bees (Bombus terrestris). J Insect Physiol (doi:10.1016/j.jinsphys.2017.11.006)

22. Sepúlveda-Rodríguez G, Roberts KT, Araújo P, Lehmann P, Baird E. 2024 Bumblebee thermoregulation at increasing temperatures is affected by behavioral state. J Therm Biol 121, 103830. (doi:10.1016/J.JTHERBIO.2024.103830)

23. McCallum KP, McDougall FO, Seymour RS. 2013 A review of the energetics of pollination biology. Journal of Comparative Physiology B 183:7 183, 867–876. (doi:10.1007/S00360-013-0760-5)

24. Heinrich B. 1975 Thermoregulation in bumblebees - II. Energetics of warm-up and free flight. Journal of Comparative Physiology B (doi:10.1007/BF00706595)

25. Kenna D, Cooley H, Pretelli I, Ramos Rodrigues A, Gill SD, Gill RJ. 2019 Pesticide exposure affects flight dynamics and reduces flight endurance in bumblebees. Ecol Evol (doi:10.1002/ece3.5143)

26. Gill RJ, Raine NE. 2014 Chronic impairment of bumblebee natural foraging behaviour induced by sublethal pesticide exposure. Funct Ecol (doi:10.1111/1365-2435.12292)

27. Kolano PJ, Borgå K, Nielsen A. 2021 Temperature sensitive effects of the neonicotinoid clothianidin on bumblebee (Bombus terrestris) foraging behaviour. J Pollinat Ecol 28, 138–152. (doi:10.26786/1920-7603(2021)633/REFERENCES)

28. Weidenmüller A, Meltzer A, Neupert S, Schwarz A, Kleineidam C. 2022 Glyphosate impairs collective thermoregulation in bumblebees. Science 376, 1122–1126. (doi:10.1126/SCIENCE.ABF7482;SUBPAGE:STRING:FULL)

29. Colgan TJ, Fletcher IK, Arce AN, Gill RJ, Ramos Rodrigues A, Stolle E, Chittka L, Wurm Y. 2019 Caste- and pesticide-specific effects of neonicotinoid pesticide exposure on gene expression in bumblebees. Mol Ecol (doi:10.1111/mec.15047)

30. Palmer MJ, Moffat C, Saranzewa N, Harvey J, Wright GA, Connolly CN. 2013 Cholinergic pesticides cause mushroom body neuronal inactivation in honeybees. Nat Commun 4, 1634. (doi:10.1038/ncomms2648)

31. Moffat C, Pacheco JG, Sharp S, Samson AJ, Bollan KA, Huang J, Buckland ST, Connolly CN. 2015 Chronic exposure to neonicotinoids increases neuronal vulnerability to mitochondrial dysfunction in the bumblebee (Bombus terrestris). FASEB Journal (doi:10.1096/fj.14-267179)

32. Mänd M, Karise R. 2015 Recent insights into sublethal effects of pesticides on insect respiratory physiology. Open access insect physiol, 31. (doi:10.2147/OAIP.S68870)

33. Lu C, Chang CH, Lemos B, Zhang Q, MacIntosh D. 2020 Mitochondrial Dysfunction: A Plausible Pathway for Honeybee Colony Collapse Disorder (CCD). Environ Sci Technol Lett 7, 254–258. (doi:10.1021/ACS.ESTLETT.0C00070/SUPPL_FILE/EZ0C00070_SI_001.PDF)

34. Brühl CA, Bakanov N, Köthe S, Eichler L, Sorg M, Hörren T, Mühlethaler R, Meinel G, Lehmann GUC. 2021 Direct pesticide exposure of insects in nature conservation areas in Germany. Scientific Reports 2021 11:1 11, 24144-. (doi:10.1038/s41598-021-03366-w)

35. Mitchell EAD, Mulhauser B, Mulot M, Mutabazi A, Glauser G, Aebi A. 2017 A worldwide survey of neonicotinoids in honey. Science 358, 109–111. (doi:10.1126/science.aan3684)

36. Zhang C, Wang X, Kaur P, Gan J. 2023 A critical review on the accumulation of neonicotinoid insecticides in pollen and nectar: Influencing factors and implications for pollinator exposure. Science of The Total Environment 899, 165670. (doi:10.1016/J.SCITOTENV.2023.165670)

37. Gkotsi D, Begum R, Salt T, Lascaratos G, Hogg C, Chau KY, Schapira AHV, Jeffery G. 2014 Recharging mitochondrial batteries in old eyes. Near infra-red increases ATP. Exp Eye Res 122, 50–53. (doi:10.1016/J.EXER.2014.02.023)

38. Quirk BJ, Sannagowdara K, Buchmann E V., Jensen ES, Gregg DC, Whelan HT. 2016 Effect of near-infrared light on in vitro cellular ATP production of osteoblasts and fibroblasts and on fracture healing with intramedullary fixation. J Clin Orthop Trauma 7, 234–241. (doi:10.1016/j.jcot.2016.02.009)

39. Giuliani A, Lorenzini L, Gallamini M, Massella A, Giardino L, Calzà L. 2009 Low infra red laser light irradiation on cultured neural cells: effects on mitochondria and cell viability after oxidative stress. BMC Complementary and Alternative Medicine 9:1 9, 8-. (doi:10.1186/1472-6882-9-8)

40. Albarracin R, Eells J, Valter K. 2011 Photobiomodulation Protects the Retina from Light-Induced Photoreceptor Degeneration. Invest Ophthalmol Vis Sci 52, 3582–3592. (doi:10.1167/IOVS.10-6664)

41. Rojas JC, Lee J, John JM, Gonzalez-Lima F. 2008 Neuroprotective Effects of Near-Infrared Light in an In Vivo Model of Mitochondrial Optic Neuropathy. Journal of Neuroscience 28, 13511–13521. (doi:10.1523/JNEUROSCI.3457-08.2008)

42. Eells JT et al. 2004 Mitochondrial signal transduction in accelerated wound and retinal healing by near-infrared light therapy. Mitochondrion 4, 559–567. (doi:10.1016/J.MITO.2004.07.033)

43. Weinrich TW, Coyne A, Salt TE, Hogg C, Jeffery G. 2017 Improving mitochondrial function significantly reduces metabolic, visual, motor and cognitive decline in aged Drosophila melanogaster. Neurobiol Aging 60, 34–43. (doi:10.1016/J.NEUROBIOLAGING.2017.08.016)

44. Begum R, Calaza K, Kam JH, Salt TE, Hogg C, Jeffery G. 2015 Near-infrared light increases ATP, extends lifespan and improves mobility in aged Drosophila melanogaster. Biol Lett 11. (doi:10.1098/RSBL.2015.0073/62394)

45. Powner MB, Priestley G, Hogg C, Jeffery G. 2021 Improved mitochondrial function corrects immunodeficiency and impaired respiration in neonicotinoid exposed bumblebees. PLoS One 16, e0256581. (doi:10.1371/JOURNAL.PONE.0256581)

46. Michael BP, Thomas ES, Chris H, Glen J. 2016 Improving Mitochondrial Function Protects Bumblebees from Neonicotinoid Pesticides. PLoS One 11, e0166531. (doi:10.1371/JOURNAL.PONE.0166531)

47. Andreo L, Mesquita-Ferrari RA, Grenho L, Gomes P de S, Bussadori SK, Fernandes KPS, Fernandes MH. 2021 Effects of 660-nm and 780-nm Laser Therapy on ST88-14 Schwann Cells. Photochem Photobiol 97, 198–204. (doi:10.1111/PHP.13323;WGROUP:STRING:PUBLICATION)

48. Softić A, Cipurković S, Avdić A. 2024 Effects of 660 nm red light and 850 nm near infrared light on human blood lymphocytes. J Phys Conf Ser 2930, 012005. (doi:10.1088/1742-6596/2930/1/012005)

49. Wickham H. 2016 ggplot2: Elegant graphics for data analysis. Springer-Verlag New York. See https://ggplot2.tidyverse.org.

50. Lüdecke D, Ben-Shachar MS, Patil I, Waggoner P, Makowski D. 2021 performance: An R Package for Assessment, Comparison and Testing of Statistical Models. J Open Source Softw 6, 3139. (doi:10.21105/JOSS.03139)

51. Kuo Y, Lu YH, Lin YH, Lin YC, Wu YL. 2023 Elevated temperature affects energy metabolism and behavior of bumblebees. Insect Biochem Mol Biol 155. (doi:10.1016/j.ibmb.2023.103932)

52. Moffatt L, Núñez JA. 1997 Oxygen consumption in the foraging honeybee depends on the reward rate at the food source. J Comp Physiol B 167, 36–42. (doi:10.1007/S003600050045/METRICS)

53. Sargent C, Ebanks B, Hardy ICW, Davies TGE, Chakrabarti L, Stöger R. 2021 Acute Imidacloprid Exposure Alters Mitochondrial Function in Bumblebee Flight Muscle and Brain. Frontiers in Insect Science 1, 765179. (doi:10.3389/FINSC.2021.765179/BIBTEX)

54. Williamson SM, Willis SJ, Wright GA. 2014 Exposure to neonicotinoids influences the motor function of adult worker honeybees. Ecotoxicology (doi:10.1007/s10646-014-1283-x)

55. Muth F, Leonard AS. 2019 A neonicotinoid pesticide impairs foraging, but not learning, in free-flying bumblebees. Sci Rep 9, 4764.

56. Lämsä J, Kuusela E, Tuomi J, Juntunen S, Watts PC. 2018 Low dose of neonicotinoid insecticide reduces foraging motivation of bumblebees. Proceedings of the Royal Society B: Biological Sciences 285. (doi:10.1098/RSPB.2018.0506/84717)

57. Suarez RK. 2000 Energy Metabolism during Insect Flight: Biochemical Design and Physiological Performance Physiological and biochemical zoology, 73(6), 765–771.. 10.1086/318112 73, 765–771. (doi:10.1086/318112)

58. Laycock I, Cotterell KC, O’Shea-Wheller TA, Cresswell JE. 2014 Effects of the neonicotinoid pesticide thiamethoxam at field-realistic levels on microcolonies of Bombus terrestris worker bumble bees. Ecotoxicol Environ Saf 100, 153–158. (doi:10.1016/j.ecoenv.2013.10.027)

59. Nieh JC, León A, Cameron S, Vandame R. 2006 Hot bumble bees at good food: thoracic temperature of feeding Bombus wilmattae foragers is tuned to sugar concentration. J Exp Biol 209, 4185–4192. (doi:10.1242/JEB.02528)

60. Tosi S, Nieh JC, Sgolastra F, Cabbri R, Medrzycki P. 2017 Neonicotinoid pesticides and nutritional stress synergistically reduce survival in honey bees. Proceedings of the Royal Society B: Biological Sciences (doi:10.1098/rspb.2017.1711)

61. Woodcock BA, Isaac NJB, Bullock JM, Roy DB, Garthwaite DG, Crowe A, Pywell RF. 2016 Impacts of neonicotinoid use on long-term population changes in wild bees in England. Nat Commun 7, 12459. (doi:10.1038/ncomms12459)

62. Guzman LM, Elle E, Morandin LA, Cobb NS, Chesshire PR, McCabe LM, Hughes A, Orr M, M’Gonigle LK. 2024 Impact of pesticide use on wild bee distributions across the United States. Nature Sustainability 7:10 7, 1324–1334. (doi:10.1038/s41893-024-01413-8)

63. Nicholson CC et al. 2023 Pesticide use negatively affects bumble bees across European landscapes. Nature 628:8007 628, 355–358. (doi:10.1038/s41586-023-06773-3)

64. Rundlöf M et al. 2015 Seed coating with a neonicotinoid insecticide negatively affects wild bees. Nature 521, 77–U162. (doi:10.1038/nature14420)

65. Tsvetkov N, Samson-Robert O, Sood K, Patel HS, Malena DA, Gajiwala PH, Maciukiewicz P, Fournier V, Zayed A. 2017 Chronic exposure to neonicotinoids reduces honey bee health near corn crops. Science 356, 1395–1397. (doi:10.1126/science.aam7470)

